# Human airway model reveals host-pathogen co-infection dynamics of *Staphylococcus aureus* and *Pseudomonas aeruginosa*

**DOI:** 10.64898/2026.03.08.710358

**Authors:** Katrine Madsen, Signe Lolle, Martin Saxtorph Bojer, Hanne Ingmer, Helle Krogh Johansen, Søren Molin, Ruggero La Rosa

**Author notes:** **Correspondence**: Ruggero La Rosa.

## Abstract

Polymicrobial airway infections, particularly those involving *Staphylococcus aureus* and *Pseudomonas aeruginosa*, are a hallmark of severe chronic respiratory diseases such as those in people with cystic fibrosis (pwCF). Yet, the mechanisms enabling their co-existence remain poorly understood. Using a human air-liquid interface (ALI) airway model, we dissected host-pathogen and interspecies dynamics during mono– and co-infections with clinically relevant bacterial pathogens. We found that *S. aureus* strains exhibited distinct colonization patterns and virulence profiles, with the deletion of the *agr* quorum sensing locus leading to attenuated epithelial disruption. Additionally, the H1 component of *P. aeruginosa*’s type VI secretion system was critical for competitive exclusion of *S. aureus,* its deletion allowing for occasional co-localization of both species. Importantly, co-infection timing strongly influenced pathogen dominance and epithelial damage, with pre-colonization of ALI models with *S. aureus* prior to *P. aeruginosa* infection, mimicking clinical observations in CF. Our results reveal that strain-specific traits and secretion systems influence microbial competition and persistence in the airway and underscore the importance of physiologically relevant models for studying chronic infections. Moreover, they provide the foundation for exploring therapeutic strategies for polymicrobial infections that account for the colonization sequence and microbial evolution for treatment design.

## INTRODUCTION

Multi-species microbial airway infections can cause severe exacerbation of pulmonary diseases, especially in people with cystic fibrosis (pwCF)^1–4^. This disease is characterized by dehydration of the airway secretion and excess mucus production, resulting in a microenvironment favouring the presence of certain bacterial species, including, but not limited to, *Staphylococcus aureus* and *Pseudomonas aeruginosa*^5–9^. *S. aureus* is among the first pathogens to colonize the airways of young children with CF^10^. In most cases, *P. aeruginosa*, which is considered a later colonizer, displaces and outcompetes *S. aureus,* becoming the dominating airway pathogen^10^. This inverse correlation between *S. aureus* and *P. aeruginosa* during years of prolonged infection in the airways has been shown in several pwCF^11^. However, longitudinal analysis of pwCF airways infections has also shown that the two pathogens can co-exist within the same patient for extended periods of time^12^. This indicates that, albeit active competition between *S. aureus* and *P. aeruginosa* might be the primary driver of the observed anticorrelation, the two bacterial species can nonetheless co-exist in the airways of pwCF, through mechanisms that remain poorly understood^13^.

Competition between *S. aureus* and *P. aeruginosa* has been extensively studied in laboratory conditions *in vitro* using both plates and liquid cultures, which do not resemble the *in vivo* conditions of the host^14–16^. Hence, limited knowledge is available on the molecular mechanisms enabling their co-existence *in vivo*. Importantly, *S. aureus* and *P. aeruginosa* co-existence severely affects patient morbidity and mortality, which is of high clinical relevance^12^. Understanding of their colonization and infection dynamics, and the interspecies interactions within the host during infection, is, therefore, of high priority to optimize treatment and improve patients’ outcomes^4^.

To establish a long-term persistent infection, *S. aureus* often acquires mutations on the *agr* operon^17,18^, influencing disease progression^19,20^. *agr* is the primary quorum sensing system in *S. aureus,* and it constitutes a major regulator of virulence, including the expression of a wide array of secreted toxins, enzymes, and quorum sensing molecules controlling toxicity towards eukaryotic cells^21^. Its downregulation is often associated with a shift towards a more persistent, less immunogenic phenotype that favours long-term colonization over acute infection^17,22,23^. Similarly, years of within-patient evolution selects for *P. aeruginosa* strains with reduced virulence, slow growth, and immune evasion, that facilitate the establishment and maintenance of a persistent infection^24–27^. *In vitro* competition studies have revealed the importance of the type VI secretion system (T6SS), specifically the H1-Type-VI-Secretion-System (H1-T6SS) of *P. aeruginosa,* in mediating antagonistic interactions against co-existing bacterial species^28^. The H1-T6SS system delivers toxins such as the Tse1-Tse7 into neighboring bacterial cells, hereby disrupting and degrading membranes and cell walls of competing bacteria, along with interfering with their replication and growth^29–32^. Recently, it was shown that different isolates of *P. aeruginosa* sampled from the same individual at a single timepoint expressed different T6SS effectors. However, the number of effector genes did not influence the possibility that a strain could cause chronic infection^33^. By eliminating competing microbes, the T6SS provides *P. aeruginosa* a significant fitness advantage in multispecies communities, being an important weapon for dominating the ecological niche^16,34,35^. Increased activity of the T6SS might be one of the causes of the described *S. aureus* and *P. aeruginosa* anticorrelation.

Recently, the development of new advanced infection model systems, such as the air-liquid interface (ALI) system, which recreates the physiology of the airway environment^36^, has allowed the study of both *S. aureus* and *P. aeruginosa* infections in host-like conditions^25,26,36–38^. However, their precise role in airway colonization during co-infection within the human host remain to be determined. Furthermore, the extent to which colonization patterns and interspecies microbial competition are strain-specific is still an open question. Addressing these points requires model systems that accurately recreate the host infection’s microenvironment and by employing clinically relevant microbial variants selected through within-host evolution. Importantly, understanding these interactions is not only critical for CF but may also be valuable for other chronic respiratory diseases characterized by polymicrobial infections.

Here, we investigated host-microbe interactions using an ALI model of human airway epithelium during infections with multiple *S. aureus* strains, including an *agr* mutant strain. This provided insight into strain-specific colonization patterns and specific host responses. Additionally, by co-infecting airway ALI models with both *S. aureus* and *P. aeruginosa,* we established a dual-species infection model system to investigate their competition and co-existence dynamics under *in vivo*-like conditions. Finally, we characterized the role of the *P. aeruginosa* H1-T6SS system in mediating co-existence and competition with *S. aureus* and its contribution to airway colonization. Altogether, our work sheds light on the infection capabilities of *S. aureus* and *P. aeruginosa* in polymicrobial infections, in a model system that closely mimics the human airway epithelium.

## RESULTS

### Strain specific *S. aureus* infection patterns in ALI models

To investigate potential differences in the colonization patterns of distinct *S. aureus* strains in conditions that mimic the human lung epithelium, we performed infection assays in ALI models based on the BCi-NS1.1 cell line. These models differentiate into a human airway epithelium representative of the *in vivo* cell composition and physiological conditions, with the apical side exposed to air and the basolateral side containing the growth medium^39^. We performed colonization studies employing the *S. aureus* strains Newman, USA300(AH1263)^40^ and JE2^41^. The Newman strain has been isolated from a human infection and shows a robust virulence phenotype^42^. The USA300(AH1263) is a hypervirulent strain due to high secretion of α-toxins known to damage epithelial tight-junctions, increased *agr-*response, and production of QS-regulated toxins^43^. Because of its widespread use in laboratories and available tools for mutagenesis, we also included the USA300 derivative JE2 strain, which exhibit a hypervirulent phenotype similar to USA300(AH1263)^43,44^.

To determine the systemic response of the host tissue, we monitored three tissue-specific indicators of the infections: transepithelial electrical resistance (TEER), lactate dehydrogenase (LDH) release, and interleukin-8 (IL-8) secretion. TEER quantifies the epithelial layer integrity, and its decrease upon infection indicates epithelial disruption. The release of lactate dehydrogenase (LDH) indicates damage of the epithelial plasma membranes and quantifies cellular cytotoxicity of cells undergoing damage and death. IL-8 is a proinflammatory cytokine released by epithelial cells upon infection, which plays an important role in immune cell recruitment independently of their presence/absence in the ALI models.

To evaluate the dynamics of host colonization, we compared changes in TEER at 16 and 24 hours post infection (hpi) relative to baseline (0 hpi). Overall, TEER decreased gradually over time for all strains consistent with their virulence profile (Fig. 1A). At 16 hpi, the Newman strain showed only minor reduction in TEER, while the USA300 and JE2 strains, showed a more pronounced decrease with TEER values already dropping to around 0.6 and 0.7, respectively. At 24 hpi, TEER decrease was substantial for all strains (Newman = 0.39 ± 0.070; JE2 = 0.26 ± 0.017), with the USA300 strain exhibiting the greatest TEER drop (0.14 ± 0.038) (Fig. 1A). For all strains, we observed an increase in LDH release at 24 hpi compared to 16 hpi (Two-way Anova with Tukey’s post hoc test, p<0.01). Moreover at 24 hpi, the USA300 strain caused significantly higher LDH release relative to both Newman and JE2 infections (Two-way Anova with Tukey’s post hoc test, p<0.01) (Fig. 1B). Similarly, we observed an increase in IL-8 secretion at 24 hpi compared to 16 hpi (two-way Anova with Tukey’s post hoc test, p<0.0001), confirming a strong immune response of the epithelium upon infection (Fig. 1C). However, no differences were observed between strains. Bacterial growth (CFU/mL) was quantified at 16 hpi and 24 hpi in the apical and basolateral compartments of the ALI model, as well as from bacteria attached to the epithelium^45^. At 16 hpi, strain showed no differences across compartments (two-way Anova with Tukey’s post hoc test, p>0.05), and no CFUs were detected basolaterally, indicating no bacterial epithelial penetration (Fig. 1D). At 24 hpi, significantly more USA300 CFUs were measured in the apical compartment compared to both Newman and JE2 (two-way Anova with Tukey’s post hoc test, p<0.0001). Both USA300 and JE2 strains were found attached to the epithelium at significantly higher CFUs compared to the Newman strain (two-way Anova with Tukey’s post hoc test, p<0.05). At 24 hpi, no bacteria were detected in the basolateral compartment following infection with the Newman strain. In contrast, basolateral bacterial counts reached 10⁵ CFU/mL for USA300 and 10³ CFU/mL for JE2, indicating strain-dependent epithelial penetration and confirming the hypervirulent phenotype of strains USA300 and JE2 (Fig. 1D). Confocal microscopy analysis revealed that at 16 hpi, all strains exhibited a similar colonization pattern with bacteria mostly dispersed as single cells across the epithelial surface (Fig. 1E). However, at 24 hpi, the USA300 and JE2 were found dispersed across the epithelium, whereas the Newman strain formed localized bigger colonies on top of the epithelium (Fig. 1E).

**Figure 1.**
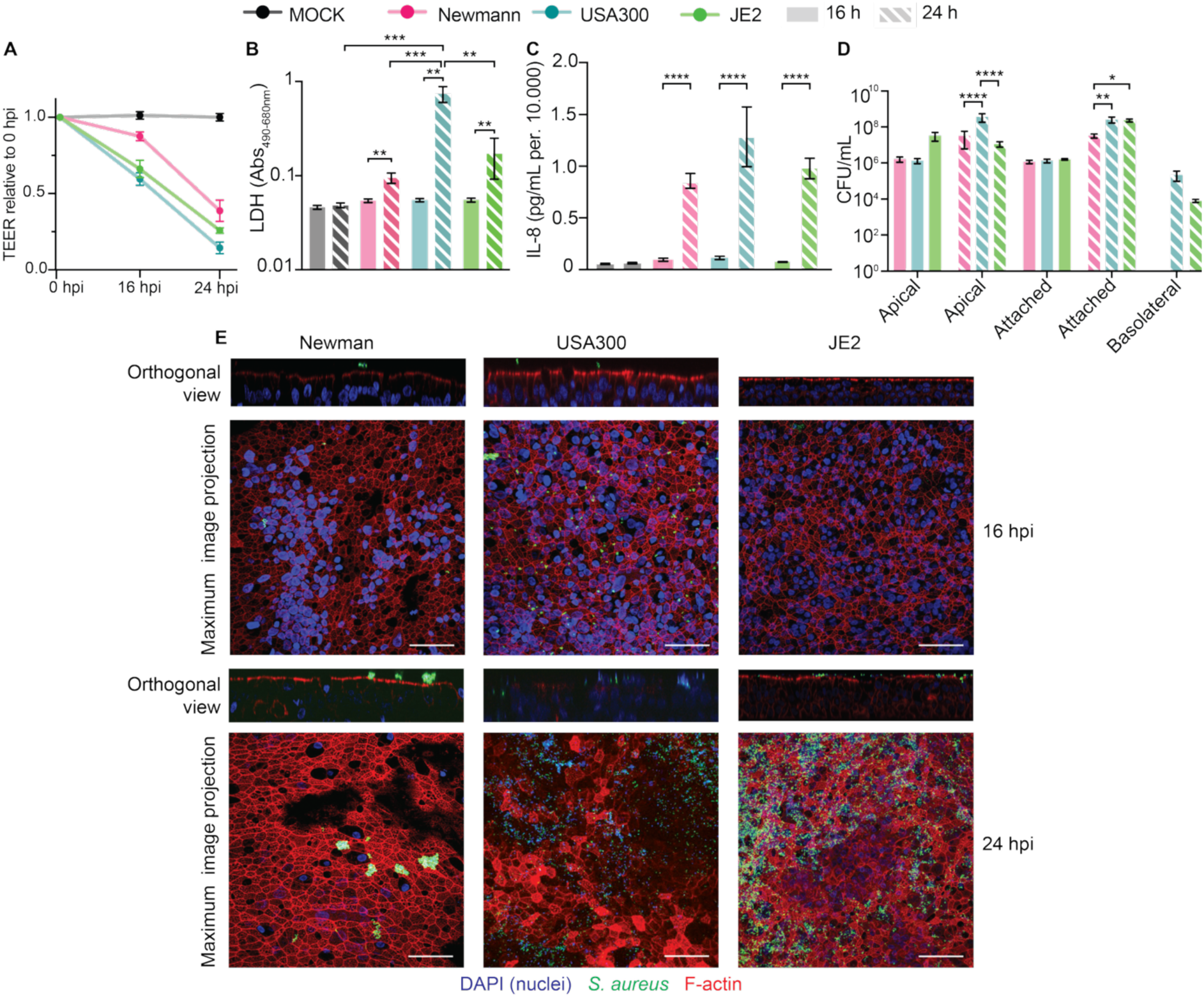
Strain specific *Staphylococcus aureus* infection patterns in air-liquid interface (ALI) infection models. **(A)** Transepithelial electrical resistance (TEER) at 16 hours post infection (hpi) and 24 hpi relative to 0 hpi. (**B)** LDH release measured as Abs_490-680_. **(C)** IL-8 release measured in pg/mL. **(D)** Bacterial growth (CFU/mL) in the apical and basolateral compartment and attached to the epithelium. Infections were performed at 16 and 24 hours post infection (hpi); unless noted, 16 hpi is shown with solid bars and 24 hpi with dashed bars. In grey mock uninfected epithelial cells, in pink Newman strain, in blue USA300 strain, and in green JE2 strain. **(E)** Confocal microscopy images of USA300, Newman and JE2 infections at 16 hpi and 24 hpi. In blue (DAPI) human nuclei, in red (phalloidin RFP) F-actin and in green (GFP) *S. aureus*. Images were acquired at 40 X and scale bar = 50 μm. In all cases, data are shown as mean ± standard error of the mean (SEM) of three independent biological replicates. Statistical significance was determined by two-way Anova with Tukey’s post hoc test where *p<0.05, **p<0.01, ***p<0.001, ****p<0.0001.

These results indicate that *S. aureus* strains exhibit distinct colonization patterns and host interaction profiles in ALI models, highlighting the importance of strain-specific characteristics in epithelial disruption, immune activation, and tissue invasion.

### *agr* mutation suppresses *S. aureus* virulence towards the airway epithelium

Agr is a major regulator of quorum sensing and virulence in *S. aureus*^22,23^, and we hypothesized a change in infection dynamics as a consequence of its disruption. For this, we investigated the colonization capacity of the JE2 strain and its isogenic *Δagr* mutant strain in ALI infection models.

At 24 hpi, the JE2 strain caused a TEER decline to around 0.3-fold, whereas the TEER remained largely unchanged for the *Δagr* infections compared to 0 hpi (one-way Anova with Tukey’s post hoc test, p<0.0001) (Fig. 2A). This lower degree of disruption of the epithelial layer by the *Δagr* mutant was further supported by LDH measurements, where the JE2 infections caused a substantially higher release of LDH compared to infection with the *Δagr* strain and mock (one-way Anova with Tukey’s post hoc test, p<0.05) (Fig. 2B). No significant difference in IL-8 release was observed between JE2 and *Δagr* strains indicating bacterial recognition by the epithelium despite the mutation (one-way Anova with Tukey’s post hoc test, p<0.0001) (Fig. 2C). Equal number of CFU/mL were found in the apical side, whereas significantly more JE2 were found attached to the epithelium compared to the *Δagr* (two-way Anova with Tukey’s post hoc test, p<0.0001) (Fig. 2D). Notably, only the JE2 strain was able to penetrate the epithelial layer as evidenced by its presence in the basolateral compartment (Fig. 2D). Confocal microscopy analyses showed that at 24 hpi, the JE2 strain was scattered on top of the epithelial layer (Fig. 2E). On the contrary, the *Δagr* strain was localized in condensed colonies on top of the epithelium, and no penetration was observed (Fig. 2E).

**Figure 2.**
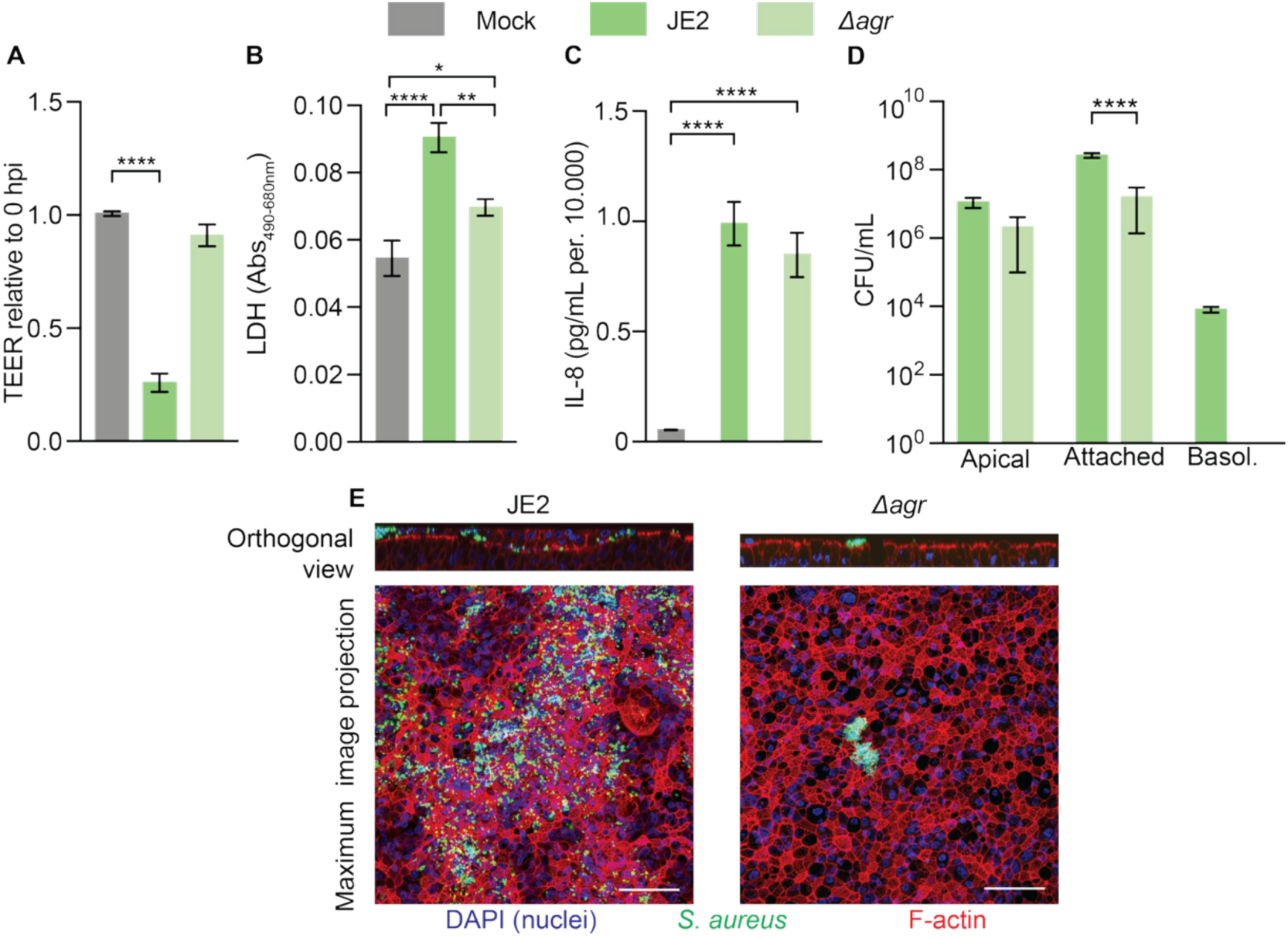
Effects of *agr* mutation on *Staphylococcus aureus* colonization and virulence. (**A)** Transepithelial electrical resistance (TEER) of mock (grey), JE2 (green) and *Δagr* (light green) at 24 hours post infection (hpi) relative to 0 hpi. **(B)** LDH release measured as Abs_490-680_. **(C)** IL-8 release measured in pg/mL. **(D)** Bacterial growth (CFU/mL) in the apical and basolateral compartment and attached to the epithelium. **(E)** Confocal microscopy images of JE2 and *Δagr* at 24 hpi. In blue (DAPI) human nuclei, in red (phalloidin RFP) F-actin and in green (GFP) *S. aureus*. Images were acquired at 40 X and scale bar = 50 μm. In all cases, data are shown as mean ± standard error of the mean (SEM) of three independent biological replicates. Statistical significance was determined by one-way (A, B, C) or two-way (D) Anova with Tukey’s post hoc test where *p<0.05, **p<0.01, ****p<0.0001.

These results confirm that disruption of the *agr* system significantly attenuates *S. aureus* virulence in ALI models, leading to reduced epithelial damage, limited tissue invasion, and altered colonization dynamics.

### Impact of H1-T6SS deletion on virulence and epithelial interaction in ALI Infection models

The T6SS is a contact-dependent structure used by *P. aeruginosa* to inject effector proteins into neighboring cells, contributing to host cell interactions and interbacterial competition^35^. We analyzed the infection capabilities of a H1-T6SS deletion strain as it is the primary system responsible for interspecies antagonism, in contrast to H2– and H3-T6SS, which are involved in host cell invasion and interactions^46^.

We infected ALI models with *P. aeruginosa* PAO1 wild type and an isogenic ΔH1-T6SS derivative^47^, and evaluated host responses at 14 hpi in absence of other bacterial competitors. Both strains caused a significant drop in TEER (Fig. 3A), and increased LDH release (Fig. 3B) compared to the mock with no significant differences between the strains (one-way Anova with Tukey’s post hoc test, p<0.001). Infections with both PAO1 and ΔH1-T6SS elicited an immune response with IL-8 secretion slightly higher for PAO1 relative to ΔH1-T6SS (one-way Anova with Tukey’s post hoc test, p<0.05; Fig. 3C). Comparing the CFU/mL between the two strains did not reveal any significant differences in the apical and attached compartments (one-way Anova with Tukey’s post hoc test, p>0.05). However, we observed significantly more CFU/mL for the ΔH1-T6SS strain in the basolateral compartment relative to PAO1 (two-way Anova with Tukey’s post hoc test, p<0.001) (Fig. 3D). For both strains, confocal microscopy analysis revealed the presence of large colonies on top of the epithelium for which we did not observe any substantial differences (Fig. 3E).

**Figure 3.**
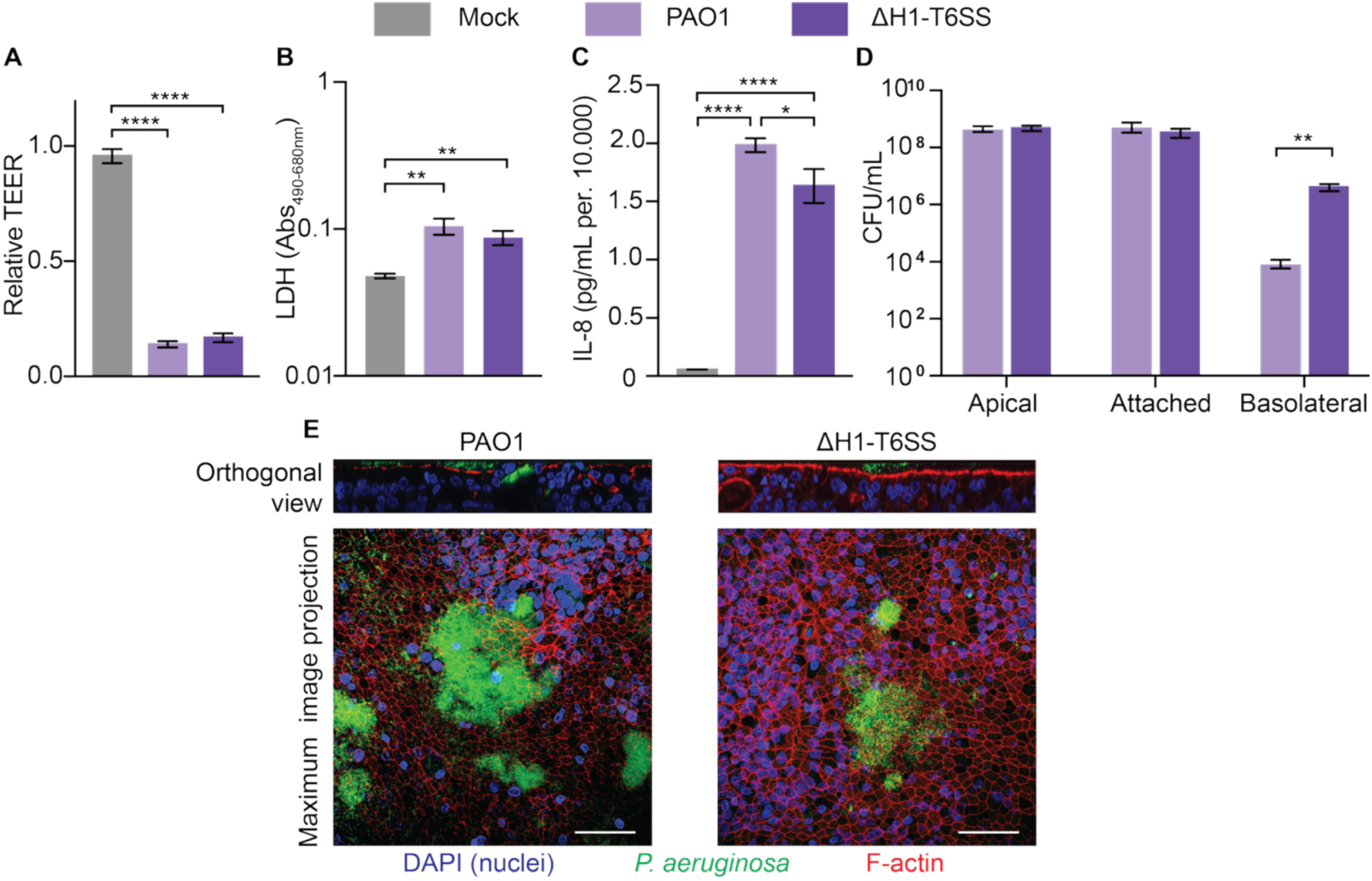
Infection phenotype of PAO1 and ΔH1-T6SS in ALI models. (**A**) Transepithelial electrical resistance (TEER) of mock (grey), PAO1 (light purple) and ΔH1-T6SS (purple) at 14 hours post infection (hpi). **(B)** LDH release measured as Abs_490-680_. **(C)** IL-8 release measured in pg/mL. **(D)** Bacterial growth (CFU/mL) in the apical and basolateral compartment and attached to the epithelium. **(E)** Confocal microscopy images of PAO1 and ΔH1-T6SS. In blue (DAPI) human nuclei, in red (phalloidin RFP) F-actin and in green (GFP) *P. aeruginosa*. Images were acquired at 40 X and scale bar = 50 μm. In all cases, data are shown as mean ± standard error of the mean (SEM) of three independent biological replicates. Statistical significance was determined by one-way (A, B, C) or two-way (D) Anova with Tukey’s post hoc test where *p<0.05, **p<0.01, ****p<0.0001.

These results suggest that deletion of the H1-T6SS does not modify epithelial damage or immune activation in ALI models but might promote epithelial penetration. Since no difference in bacterial growth rate (ΔH1-T6SS = 1.2 h^-1^ ± 0.05; PAO1 = 1.1 h^-1^ ± 0.03) was observed neither in laboratory conditions nor in ALI infection (apical and attached compartment), this result suggests a potential additional role of the H1-T6SS in modulating host interactions.

### Colonization order and H1-T6SS function modulate *S. aureus*-*P. aeruginosa* interactions and host responses in airway epithelium

To investigate how colonization timing and order influence interspecies dynamics and host responses, we performed co-infection assays in ALI models using *S. aureus* and *P. aeruginosa* either added simultaneously, as commonly done *in vitro,* or sequentially with *S. aureus* preceding *P. aeruginosa* to mimic the typical colonization pattern in pwCF.

When either USA300(AH1263) or Newman strain were added simultaneously in equal amounts with PAO1 in the ALI model, PAO1 outcompeted both strains, leaving close to no visible *S. aureus* growth at the end of the infection and only PAO1 was recovered in the ALI compartments (Fig. S1).

In contrast, when the JE2 strain was allowed to colonize the ALI model for 24 h prior to the addition of PAO1 and the co-infection was analyzed after 14 h, equal amounts of bacteria (CFU/mL) were recovered for both pathogens both in the apical compartment and attached to the epithelium (two-way Anova with Tukey’s post hoc test, p>0.05) (Fig. 4A, see JE2 (PAO1 co-culture) *vs* PAO1 (JE2 co-culture)). Only *P. aeruginosa* was recovered from the basolateral compartment (Fig. S2A), which is consistent with previous findings showing its ability to outcompete *S. aureus* in liquid cultures^48^. Similar results were obtained for co-infected cultures of either *S. aureus* Newman or USA300 with *P. aeruginosa*, underlining the importance of the order of colonization despite intrinsic differences between *S. aureus* strains (Fig. S3). Prolonged co-infection times with *S. aureus* and *P. aeruginosa* resulted in severely damaged epithelium, and no reliable quantitative measurements could be obtained. The host responses to the co-infections (TEER, LDH and IL-8) were comparable to those observed for infections with PAO1 alone and exacerbated relative to JE2 single infection (one-way Anova with Tukey’s post hoc test, p<0.05) (Fig. 4B-D). The presence of both pathogens was supported by confocal microscopy analysis showing that both *S. aureus* (in green) and *P. aeruginosa* (in red) were homogeneously distributed and co-colonized the airway epithelium (Fig. 4E).

**Figure 4.**
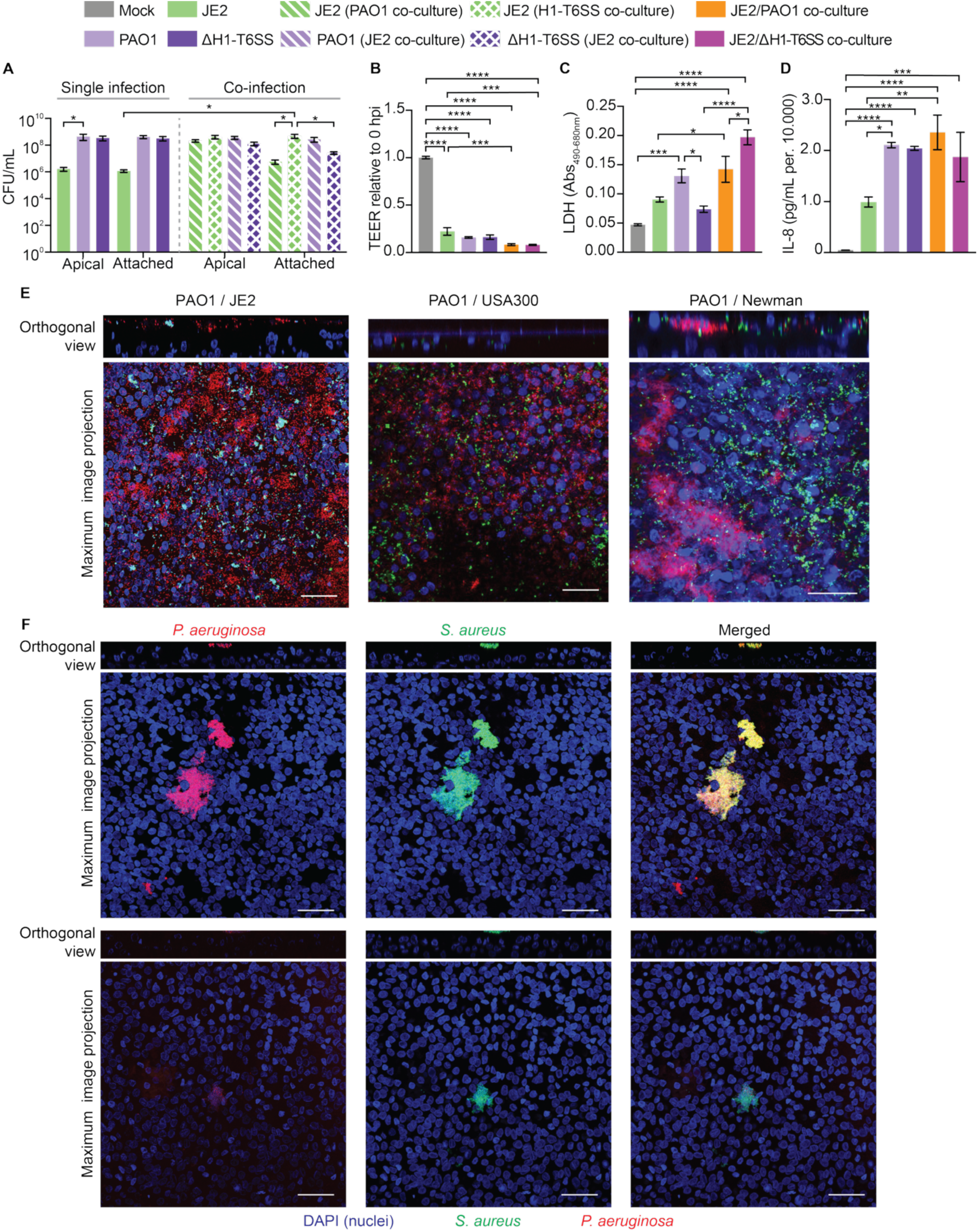
Dynamics of Staphylococcus aureus and Pseudomonas aeruginosa co-infection. (**A)** Bacterial growth (CFU/mL) in the apical and basolateral compartment and attached to the epithelium at 38 hours post infection (hpi) (24 h *S. aureus* pre-incubation plus 14 h of *P. aeruginosa* co-infection). **(B)** Transepithelial electrical resistance (TEER). **(C)** LDH release measured as Abs_490-680_. **(D)** IL-8 release measured in pg/mL. **(E-F)** Confocal microscopy images of *S. aureus* and *P. aeruginosa* in co-infection. In blue (DAPI) human nuclei, in red (RFP) *P. aeruginosa*, and in green (GFP) *S. aureus*. Images were acquired at 40 X and scale bar = 50 μm. In **(F)**, images show examples of micro-colonies of *S. aureus* and *P. aeruginosa* ΔH1-T6SS where both species are colocalized. In all cases, data are shown as mean ± standard error of the mean (SEM) of three independent biological replicates. Statistical significance was determined by one-way (B, C, D) or two-way (A) Anova with Tukey’s post hoc test where *p<0.05, **p<0.01, ***p<0.001, ****p<0.0001.

Since the T6SS of *P. aeruginosa* is directly involved in interspecies competition, we co-infected the ALI model with the JE2 strain and the ΔH1-T6SS mutant strain. Bacterial growth analysis (CFU/mL) revealed significantly more JE2 strain attached to the epithelium when co-infected with the ΔH1-T6SS mutant strains compared to PAO1 (two-way Anova with Tukey’s post hoc test, p<0.05) (Fig. 4A). TEER drop, LDH release and IL-8 secretion were, instead, comparable between co-infections with a slightly higher LDH release for ΔH1-T6SS strain co-infection (one-way Anova with Tukey’s post hoc test, p<0.05) (Fig. 4B-D). Confocal microscopy analysis showed that the colonization patterns were generally comparable to the co-infection with PAO1 and JE2 (Fig. S2B). However, thorough analysis across several samples revealed confined colonies consisting of both pathogens, suggesting the possibility of co-colony formation when *P. aeruginosa* lacks a functioning H1-T6SS (Fig. 4F and S2A).

Together, these results show that colonization order and the presence of a functional H1-T6SS partly shape the outcome of *S. aureus* and *P. aeruginosa* interactions in airway epithelium, influencing both bacterial distribution and host responses.

## DISCUSSION

Polymicrobial infections are often associated with increased morbidity and mortality in several diseases. In pwCF, *S. aureus* colonizes the airways for several years as the dominant pathogen^34^. Over time, it is typically substituted by *P. aeruginosa* driven by competitive interactions between the two species^10^. However, while clinical observations on this negative correlation are available, molecular understanding of their competition, co-existence and infection dynamics are lacking. We found that the order in which *S. aureus* and *P. aeruginosa* infect airway epithelial cells plays a critical role in determining species dominance. Indeed, early colonization may allow a pathogen to modify the epithelial surface, compete for nutrients, and trigger host responses that may inhibit or facilitate subsequent colonization by another species. For example, *P. aeruginosa* introduced first may produce antimicrobial compounds or disrupt epithelial integrity, limiting *S. aureus* colonization, whereas *S. aureus* arriving first may establish protective niches that resist displacement^49^. Clinically, this mirrors observations in pwCF, where *S. aureus* often colonizes early in life, with *P. aeruginosa* taking over in later stages^56^. *In vitro* studies have demonstrated that *P. aeruginosa* produces several virulence factors such as pyocyanin, rhamnolipids, and HQNO (2-heptyl-4-hydroxyquinoline N-oxide) that inhibit *S. aureus*. In response, *S. aureus* can adapt by switching to small colony variants, a phenotype characterized by slow growth, reduced metabolism, and increased resistance to oxidative stress and antibiotics, which facilitates persistence under such conditions^51^.

Interestingly, co-infection with the H1-T6SS mutant strain allowed for a higher *S. aureus* count during infection due to the reduced interspecies competitive advantage of *P. aeruginosa*^28–31^. Additionally, we identified some confined microcolonies where both species were present, in contrast to the otherwise dispersed colonization patterns observed with the wild type PAO1. This suggests that mutations in the H1-T6SS could facilitate the occasional formation of stable co-infection microcolonies by preventing *S. aureus* killing, antagonism and *P. aeruginosa* dominance. However, further studies are needed to further investigate the involvement of H1-T6SS within co-infected individuals.

Importantly, co-infection with *S. aureus* and *P. aeruginosa* can significantly modulate bacterial susceptibility to antibiotics by both increasing resistance or susceptibility to multiple antibiotics, as previously reported^52–54^. These results highlight the complex nature of interspecies interactions and the importance of their consideration when selecting treatment regimens for polymicrobial infections. Poor correlation between *in vitro* susceptibilities to antibiotics and the patients clinical response currently poses a challenge in the treatment of infections^55^. Therefore, testing antibiotic efficacy in multispecies infections within a human-relevant model may, indeed, help understanding and improve therapeutic strategies for treatment-resistant polymicrobial infections^25,27^. Additionally, recreating a more physiologically and clinically relevant infection model system to study both host-pathogen and pathogen-pathogen interactions could provide new knowledge on treatment failure, host responses in polymicrobial infections, and infection capabilities of bacterial strains.

Given the different virulence profiles and infection capabilities of *S. aureus* strains, appropriate infection models are fundamental to characterize infection progression and host responses. Indeed, *S. aureus* strains showed marked differences in ALI models linked to their virulence profile. Additionally, mutation of the *agr* regulator fully attenuated the virulence of *S. aureus* by changing the bacterial colonization capacity, reducing epithelial damage, and limiting immune system recruitment. This is in line with the presence of *agr*-defective isolates in patients with pneumonia^17,18,56^. Recent works suggest that elevated levels of sialic acid may confer a fitness advantage to *agr* mutants, suggesting that the reduced epithelial damage observed in ALI models may represent only one aspect of the benefits associated with *agr* dysfunction^34^. However, further investigation are required to fully characterize the adaptive advantages conferred by *agr* inactivation^57^.

Altogether these results, obtained in a controlled yet physiologically relevant *in vitro* co-infection system, pave the way for deeper exploration of interspecies bacterial dynamics and infection underlining the critical role of the human airway epithelium in shaping infection outcomes.

## METHODS

### Bacterial culture and strains

The following strains were used for the project: *S. aureus:* Newman, USA300(AH1263)^58^, JE2, JE2*(Δagr),* and *P. aeruginosa* PAO1, PAO1(ΔH1-T6SS)^47^. Strains were routinely grown at 37 °C, *S. aureus* on Tryptic Soy Broth media and *P. aeruginosa* on LB media.

### GFP and mKOk tagging of *P. aeruginosa*

*P. aeruginosa* strains were tagged either with a superfolder green fluorescent protein (sfGFP) or with mKusabira-Orange-kappa (mKOk) to enable visualizason by confocal microscopy. Bacterial strains were tagged by four parental masng using the miniTn7 delivery plasmid (pJM220)^59,60^. The plasmid was engineered to accommodate the sfGFP or mKOK gene. Transformants were selected on Pseudomonas isolason agar (PIA) supplemented with 30 µg/mL gentamicin. The final tagged strains were verified by fluorescence using a Leica DM4000 B epifluorescence microscope and growth rate was not affected by chromosomal tagging.

### GeneraXon of JE2(Δ*agr*)

The *agr* deletion strain JE2 was generated by phage transduction (phi11) of the deletion from RN6911^61^ and selecson on tetracycline (5 mg/l). The strain was verified phenotypically by lack of hemolysis on blood plates and by deficiency in autoinducing pepsde producson.

### GFP tagging of *S. aureus* strains

*Staphylococcus* strains were marked with GFP by transformation with pCM11^62^ and selection on erythromycin (10 mg/l).

### Air-liquid Interface model

BCi-NS1.1^39^ cells were seeded in Pneumacult^TM^ – ExPlus Medium (STEMCELL Technologies) and expanded in T25 culture flasks in incubator 37C, 5% CO_2_ following manufacturers’ instruction. Confluent cells with a viability >80 % were seeded (1.5*10^5^ cells/well) in collagen (Gibco) coated transwells (1.0 μm pore size; Falcon^®^). Cells were kept in Pneumacult^TM^– ExPlus Medium until confluent and subsequently airlifted. 400 µL Pneumacult^TM^ – ALI maintenance medium (STEMCELL technologies) was added to the basolateral chamber and cells were left to differentiate for 28 days, with media change twice a week. TEER was measured once a week using EVOM2^TM^ electrode. Mucus was frequently removed by PBS wash of apical compartment.

### Infection and characterization

Overnight cultures of bacteria were grown in 6 mL TSB or LB respectively in 15 mL tubes while shaking and angled 45°. OD_600_ of the ON culture was measured and diluted to OD_600_ = 0.1 (infection culture) and left shaking in incubator at 37 °C for 4-5 hours. 1 mL of infection culture was centrifuged for 5 min. at 4500 x *g,* the pellet was resuspended in D-PBS to reach OD_600_ = 1. The culture was then diluted to reach 1*10^5^ CFU/mL. 10 µL of the culture (1*10^5^ CFU/mL) was inoculated to the epithelial layer on the apical side. Transwells were incubated at 37 °C with 5 % CO_2_ for the indicated time (see individual experiments). Infection was ended by adding 200 µL PBS to the apical side and TEER was measured using EVOM2^TM^ electrode.

### CFU, LDH and IL-8 quantification

For CFU count, cells were harvested from all the apical and basolateral compartments of the ALI model plated in serial dilutions of 10 µL per droplet. Bacteria attached to the epithelium were quantified by scraping off the epithelium. *S. aureus* and *P. aeruginosa* were discriminated on agar plates based on morphology.

From the infection, basolateral media was collected and used for LDH and IL-8 analysis. LDH was measured using The Invitrogen™ CyQUANT™ LDH Cytotoxicity Assay Kit (Cat. No. C20301) following manufacturers’ instruction. IL-8 was measured using Human IL-8/CXCL8 DuoSet ELISA kit (R&D Systems).

### Confocal microscopy analysis

Cells were fixated for confocal microscopy analysis using 4 % paraformaldehyde (PFA) (in PBS). 400 µL PFA was added to the basolateral compartment and 200 µL to the apical compartment and left for 20 min. at 4C. PFA was removed, and cells washed three times in D-PBS. Next, 400 µL blocking buffer (3 % BSA, 1 % saponin, 1 % Triton-X in PBS, pH 7.4) was added to the basolateral compartment and 200 µL to the apical compartment and left for 30 min. at room temperature and washed three times with D-PBS. Primary antibodies were diluted in staining buffer (3 % BSA, 1 % saponin in PBS, pH 7.4) and 100 µL added to the apical part of the transwell and 400 µL staining buffer added to the basolateral compartment. Samples were left in the dark at room temperature for 2 h. Primary antibody and staining buffer was removed and transwells were washed three times for 5 min. in D-PBS. Antibodies included: DAPI diluted 1:500, Phalloidin 1:200 (Alexa fluor). Samples were washed 3 times for 5 minutes in D-PBS and liquid was aspirated from the wells. Transwell filters were cut out using a scalpel and transferred to a microscope slide. 8 µL VECTASHIELD® Antifade Mounting Medium (VWR, VECTH-1000) was added on top of the sample and sealed with a glass slide and nail polish. Microscopy slides were visualized with a Leica Stellaris 8 Confocal Microscope (40X, 1.3 oil) and analyzed using ImageJ software.

### Statistical analyses

GraphPad Prism 10.4.1 was used for statistical analysis. ALI data were represented as mean ± SEM. Biological replicates represent independent experiments performed with cells from different passages and technical replicates cells from same passage. Statistical comparisons used included calculated using one-way ANOVA and two-way ANOVA with Tukey’s post hoc test. p-value < 0.05 was considered statistically significant. All figures were completed in Adobe Illustrator Artwork 28.7.10. All data are provided within the manuscript as Source Data file.

### Data availability statement

The datasets generated during and/or analyzed during the current study are available within the paper in the Supplementary data 1.

## Acknowledgements

We thank the Infection Microbiology group at Rigshospitalet and The Technical University of Denmark for their work, insightful comments, and discussions. Authors thank Daniel Unterweger for the kind gift of the H1-T6SS mutant and its isogenic PAO1 strain. The Basal Cell Immortalized Non-Smoker 1.1 (BCi-NS1.1) cell line was a gift from Professor Ronald G. Cristal (Weil Cornell Medical College, New York, USA).

## Supplementary figures

**Supplementary figure 1.**
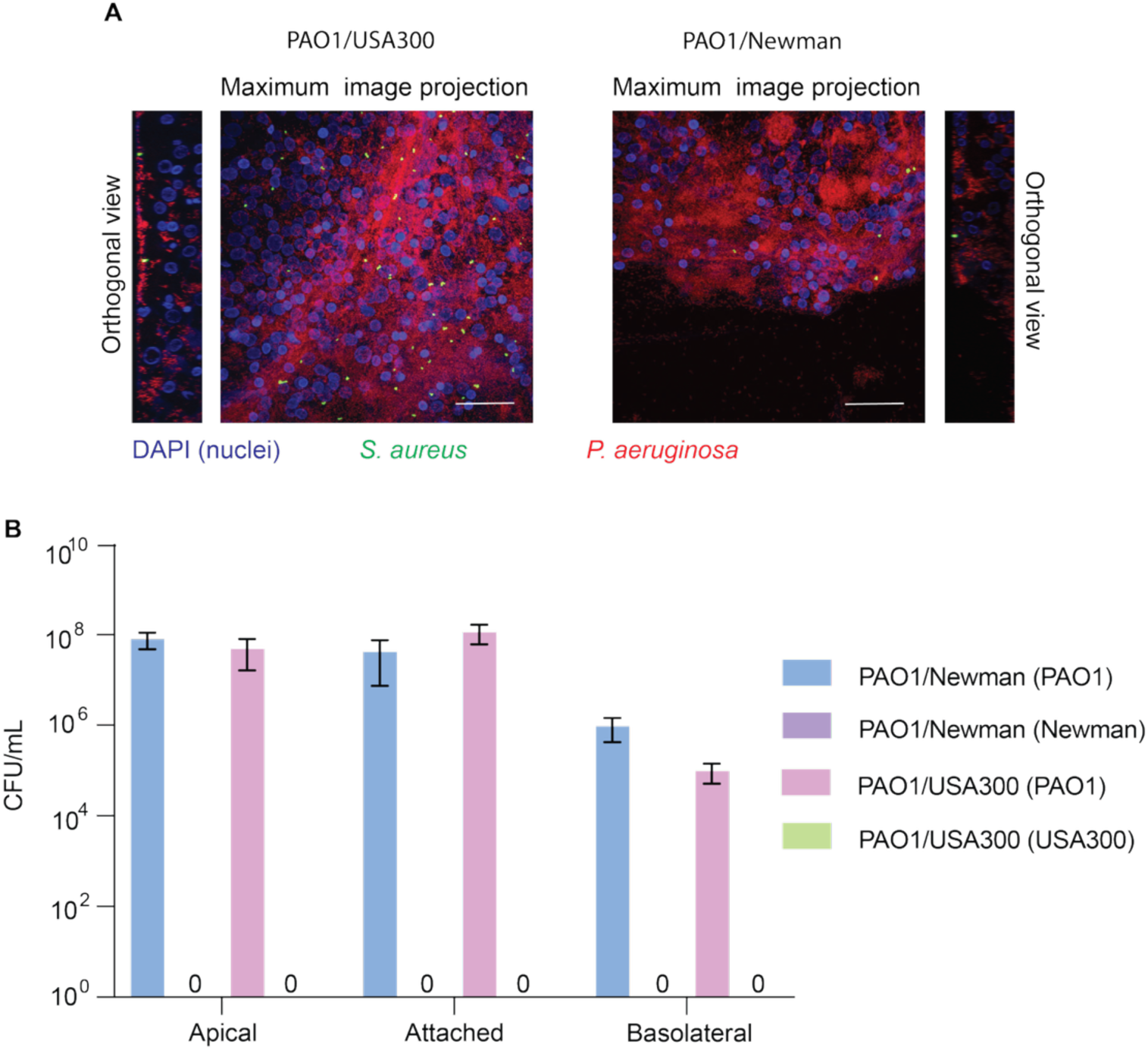
*S. aureus* and *P. aeruginosa* infection when added simultaneously to ALI models. (**A**) Confocal microscopy images of *S. aureus* and *P. aeruginosa* in co-infection when both pathogens were added in equal amounts simultaneously. *S. aureus* in green (GFP), *P. aeruginosa* in red (RFP), human nuclei in blue (DAPI). Images were acquired at 40 X and Scale bar show 50 μm. (**B**) CFU/mL in apical and basolateral compartments and attached to the epithelium. Statistical significance was determined by two-way Anova with Tukey’s post hoc test. Data are shown as mean ± standard error of the mean (SEM) of three independent biological replicates.

**Supplementary figure 2.**
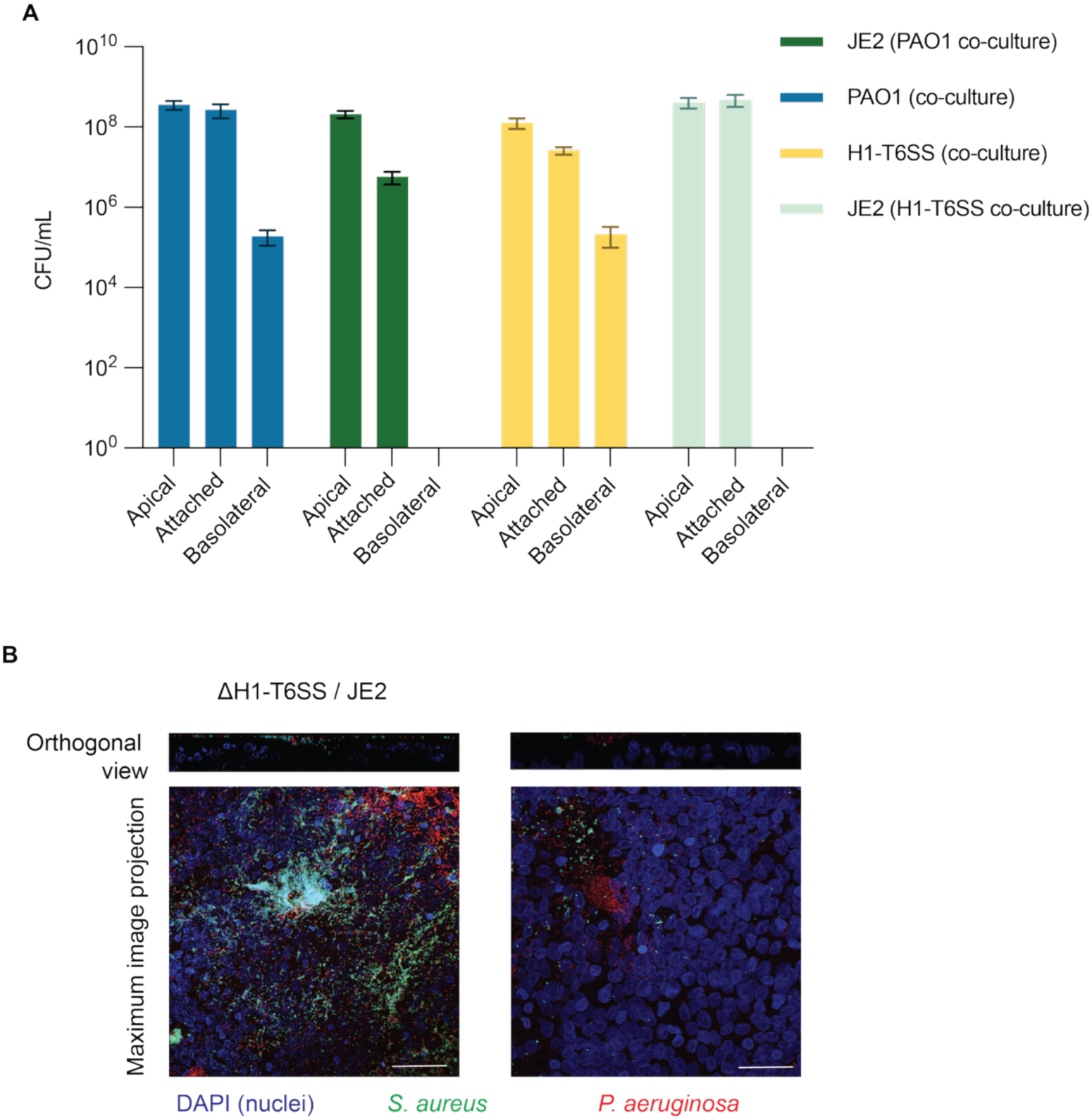
Infection parameters for JE2-PAO1 and JE2-ΔH1-T6SS infections. (**A**) CFU/mL in apical and basolateral compartments and attached to the epithelium for co-infections with PAO1/JE2 and ΔH1-T6SS/JE2. Data are shown as mean ± standard error of the mean (SEM) of three independent biological replicates. (**B**) Confocal microscopy images of co-infection with ΔH1-T6SS and JE2. *S. aureus* in green (GFP), *P. aeruginosa* in red (RFP), human nuclei in blue (DAPI). Images were acquired at 40 X and Scale bar show 50 μm.

**Supplementary figure 3.**
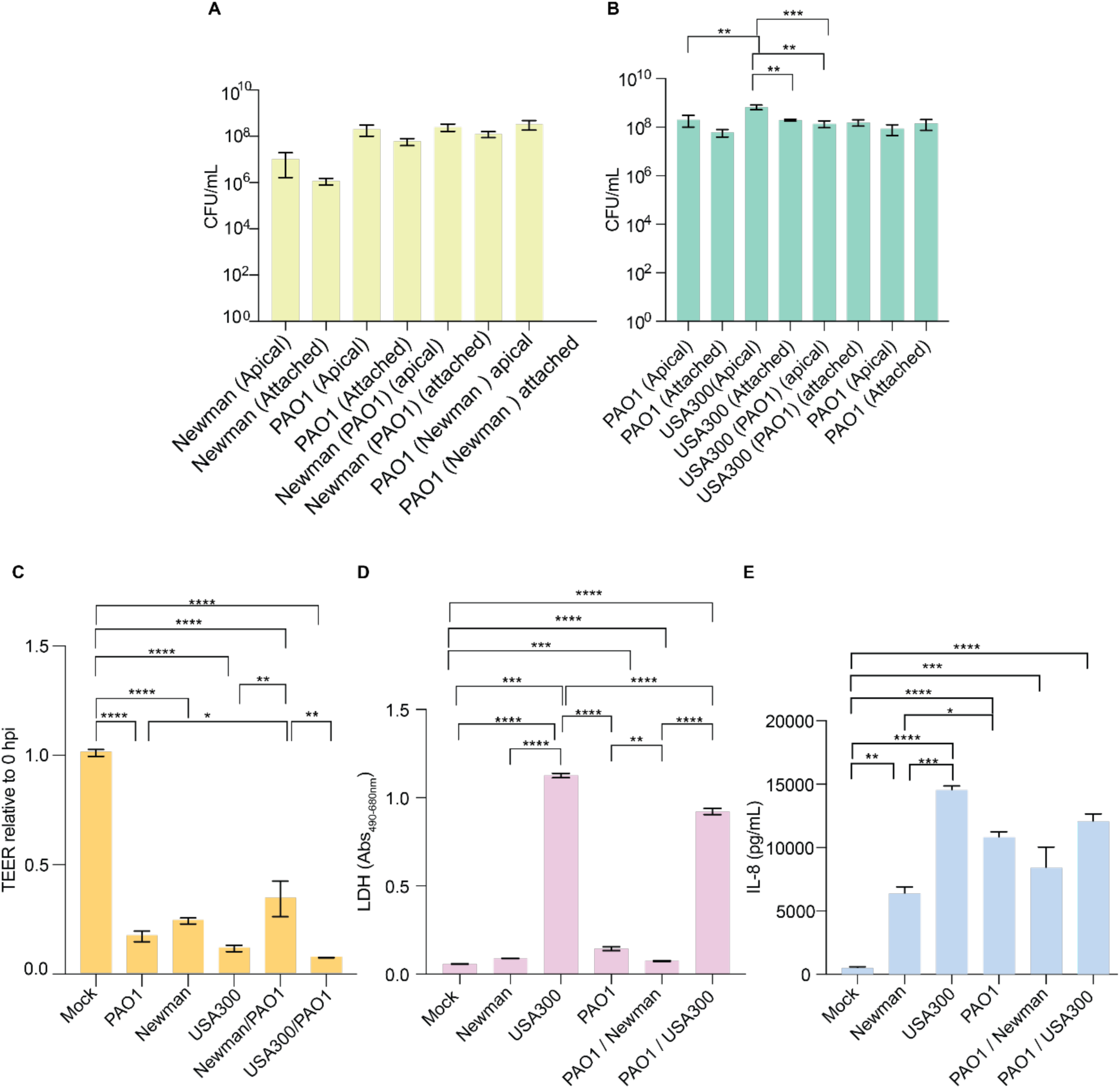
Infection parameters for Newmann-PAO1 and USA300-PAO1 con-infection. Bacterial growth (CFU/mL) in the apical and basolateral compartment and attached to the epithelium at 38 hpi (24 h *S. aureus* pre-incubation plus 14 h of *P. aeruginosa* co-infection) with strains (**A**) Newmann-PAO1 and (**B**) USA300-PAO1. Data are shown as mean ± standard error of the mean (SEM) of three independent biological replicates. Statistical significance was determined by two-way Anova with Tukey’s post hoc test where ***p<0.001, **p<0.01, *p<0.05. (**C**) Transepithelial electrical resistance (TEER) at 38 hpi. Statistical significance was determined by one-way Anova with Tukey’s post hoc test, ****p<0.0001, ***p<0.001, *p<0.05. Data are shown as mean ± standard error of the mean (SEM) of three independent biological replicates. (**D**) LDH release measured as Abs_490-680_ at 38 hpi. Statistical significance was determined by one-way Anova with Tukey’s post hoc test, ****p<0.0001, ***p<0.001, **p<0.01. Data are shown as mean ± standard error of the mean (SEM) of three independent biological replicates. (E) IL-8 release measured in pg/mL at 38 hpi. Statistical significance was determined by one-way Anova with Tukey’s post hoc test, ****p<0.0001, ***p<0.001, **p<0.01, *p<0.05. Data are shown as mean ± standard error of the mean (SEM) of three independent biological replicates.

